# Genetic architecture and evolution of stripe rust resistance uncovered using diverse panels of wheat lines and North American *Puccinia striiformis* f. sp. *tritici* isolates

**DOI:** 10.64898/2025.12.10.693471

**Authors:** Buket Sahin, Meinan Wang, Jason D. Zurn, Xiaoting Xu, Guihua Bai, Alina Akhunova, Xianming Chen, Eduard Akhunov

## Abstract

Newly emerging highly virulent races of *Puccinia striiformis* f. sp. *tritici* (*Pst*) often defeat deployed resistance genes (*Yr*), highlighting the need for novel sources of durable resistance. A global diversity panel of 377 spring wheat (*Triticum aestivum* L.) lines was screened for all-stage resistance (ASR) against 20 diverse *Pst* isolates at the seedling stage and adult plant resistance (APR) against natural mix of field races. Genome-wide association mapping identified 77 unique *Yr* loci. Of these, 33 overlapped with the previously reported ~1,100 loci, while 44 were likely novel. Nine mapped *Yr* loci were effective at the adult plant stage and showed no overlap with the known APR genes. Two wheat lines, though lacked the widely effective *Yr5* and *Yr15* genes, exhibited resistance to all 20 *Pst* races at the seedling stage and natural field races at the adult stage, suggesting that they may carry novel, broad-spectrum resistance alleles. APR genes *Yr18*, *Yr29*, and *Yr36* were detected at low frequencies, indicating that much of the observed resistance may arise from less characterized or novel sources. Wheat improvement had no effect on the frequency of ASR alleles but resulted in a three-fold increase in the frequency of APR alleles, suggesting that the latter were subjected to more consistent breeding selection over time. Our findings underscore the value of combined screening of diverse germplasm with the diverse panels of pathogen races to identify novel sources of broad-spectrum resistance for breeding stripe rust resistant cultivars.

## Introduction

Wheat (*Triticum aestivum* L.) is a cornerstone of global agriculture, cultivated on more than 220 million hectares worldwide and providing a primary source of food and nutrition for billions of people (FAOSTAT 2023). Wheat yields are consistently challenged by a variety of biotic and abiotic stress factors. Among biotic constraints, stripe rust caused by *Puccinia striiformis* f. sp. *tritici* (*Pst*) remains one of the most widespread and devastating fungal diseases affecting wheat. Over the past two decades, *Pst* races have emerged that are more virulent or adapted to warmer environments, enabling the pathogen to spread into regions previously considered unsuitable for disease development (Chen 2005). These changes have led to more frequent and severe epidemics, with annual yield losses estimated to exceed one billion US dollars (Chen 2020). The scale and persistence of these losses highlight the urgent need for cultivars that offer durable and effective resistance to evolving *Pst* populations (Beddow et al. 2015).

Genetic resistance is the most sustainable method for controlling stripe rust in wheat. Resistance is classified into two categories: all-stage resistance (ASR), which is usually race-specific and controlled by single major genes, and adult plant resistance (APR), which is governed by allelic variants of genes with small effects, providing partial but more durable protection (Singh et al. 2015; Huerta-Espino et al. 2020; Wang et al. 2023). ASR is active across all growth stages but is often overcome by new pathogen races (Ballini et al. 2013; Singh et al. 2015; Schwessinger 2016), particularly when deployed as single-gene resistance. To extend its efficacy, ASR is commonly incorporated as part of multi-gene combinations (Bulli et al. 2016; Zhang et al. 2019). In contrast, APR tends to be more stable over time due to its complex genetic basis, lower race specificity, and lower levels of resistance, which reduces selection pressure on pathogen populations (Chen 2013; Brown 2015; Singh et al. 2016). To date, 87 named *Yr* genes have been included into the wheat gene catalogue, of which 18 confer APR and the remaining provide ASR (McIntosh 2024). Modern wheat breeding programs have historically relied on elite germplasm as primary sources of resistance, largely due to their agronomic performance and adaptation (Reif et al. 2005; Lopes et al. 2015). These elite varieties carry a few multi-gene combinations selected for their ability to provide resistance to endemic *Pst* races. The effectiveness of many resistance loci in commercial varieties has declined in some regions due to the ongoing evolution of *Pst* populations (Hovmøller et al. 2016; Riella et al. 2024), including recent worrying reports of *Yr15* breakdown in Europe (Davis et al. 2025) This underscores the need to discover and utilize additional resistance sources that remain effective against newly emerging *Pst* races. Beyond elite germplasm, a wide range of genetic resources, including landraces, historical cultivars, and breeding lines, offer valuable reservoirs of disease resistance that have received limited attention in modern breeding efforts. Many of these materials were selected under local environmental pressures, allowing them to co-evolve with diverse *Pst* populations and retain greater allelic diversity (Zuev et al. 2024). In contrast to elite lines shaped by repeated selection and reduced genetic variation, older accessions often carry rare and previously untapped resistance alleles (Adhikari et al. 2012). Landraces from regions such as Turkey, Ethiopia, and Central Asia have shown both ASR and APR responses to stripe rust, underscoring their potential value in resistance breeding programs (Zegeye et al. 2014; Sehgal et al. 2016; Jia et al. 2020). However, a large proportion of accessions conserved in global gene banks remain uncharacterized for resistance to currently prevalent *Pst* races, leaving much of their genetic potential unexplored. Systematic evaluation of these underutilized resources is critical for identifying novel resistance loci and expanding the genetic base of wheat improvement.

Genome-wide association studies (GWAS) have proven to be an effective approach for dissecting the genetic architecture of complex traits, such as disease resistance, particularly when applied to germplasm collections encompassing broad natural variation (Yu et al. 2006). In wheat, GWAS has been successfully employed to detect numerous loci associated with stripe rust resistance, particularly within elite breeding materials (Kertho et al. 2015; Bulli et al. 2016; Liu et al. 2020; Gao et al. 2024; Wu et al. 2025). Nevertheless, studies incorporating genetically diverse and globally representative germplasm panels remain limited, and the extent of novel resistance variation segregating in unimproved materials is still insufficiently characterized. Expanding GWAS efforts to include these underutilized genetic resources holds substantial promise for the discovery of novel resistance loci and for enhancing the long-term efficacy of resistance breeding strategies.

Building on this premise, the present study evaluated a diverse panel of 377 spring wheat accessions from the USDA National Small Grains Collection. This panel includes landraces, historical cultivars, breeding lines, and other genetic materials representing over a century of global wheat diversity and encompassing major centers of origin, diversification, and adaptation. The primary objective was to identify genomic regions associated with resistance to stripe rust by applying genome-wide association mapping to this genetically diverse panel, with the goal of capturing both common and rare alleles that may have been beyond the resolution of prior studies focused on narrower populations. In parallel, the study aimed to dissect the genetic architecture of resistance by estimating the contribution of individual loci to phenotypic variation and by identifying genomic regions conferring consistent resistance across 20 distinct *Pst* races. This comparative, multi-race approach enables the differentiation of loci associated with ASR from those contributing to APR, supporting the prioritization of loci with stable effects across environments and pathotypes.

## Materials and methods

### Plant materials and genotyping

A genetically diverse set of 377 spring wheat accessions was selected from the USDA National Small Grains Collection to represent broad allelic variation within global wheat diversity (Table S1). The accessions include landraces, historical cultivars, breeding selections, and other genetic materials originating from 79 countries across seven geographic regions, with collection years spanning from 1904 to 2017. Selection criteria prioritized broad genetic diversity, geographic representation, and historical relevance to breeding programs.

Genotyping was performed as described by He et al. (2022), using a combined platform approach that integrated the 90K iSelect SNP array, genotyping-by-sequencing (GBS) based on restriction enzyme digestion, targeted resequencing of regulatory regions, and RNA-seq–based SNP discovery. This multi-tiered strategy provided high-density, genome-wide coverage suitable for high-resolution association analysis. Missing genotype calls were imputed with Beagle, incorporating the complete marker set (Browning et al. 2021). Quality control filtering excluded SNPs with a minor allele frequency (MAF) below 0.05, a heterozygosity rate above 3%, or missing data exceeding 50% at the marker or accession level. Following filtering, a total of 1,521,858 high-confidence SNPs was retained for downstream GWAS and genomic analyses (He et al. 2022).

In addition to genome-wide SNP genotyping, the panel was genotyped using five Kompetitive Allele-Specific Polymerase Chain Reaction (KASP) markers linked to previously characterized *Yr* resistance genes. Genotyping was performed markers available at the USDA-ARS Genotyping Laboratory in Manhattan, KS, targeting genes are *Yr5* (Marchal et al. 2018) and *Yr15* (Klymiuk et al. 2018), which confer ASR, and *Yr36* (Rasheed et al. 2016), *Lr34/Yr18* (Lagudah et al. 2009), and *Lr46/Yr29*, which are associated with APR.

### Seedling stage evaluations

Resistance to stripe rust was evaluated at the seedling stage under controlled greenhouse conditions in 2021 and 2022 using 20 *Pst* races with distinct virulence profiles (Table 1). These races were selected to represent a range of virulence combinations and included both well-established and recently identified pathotypes. Each race was phenotyped using a set of differential wheat lines carrying single *Yr* resistance genes (Wan and Chen 2014), and their virulence patterns were validated against reference isolates to ensure consistency across trials. Growing plants and inoculation were conducted following the standard procedure and conditions (Chen and Wang 2025). Briefly, 5-7 seeds of each line were planted in a pot of 6 × 6 × 6 cm filled with soil mixture containing 6 L peat moss, 2 L perlite, 3 L sand, 3 L Palouse silt loam soil, 4 L vermiculite, 250 g Osmocote 14-14-14 and 2 L water premixed in a soil mixer rotating for 30 min. The pots were placed in a greenhouse set at 10-25 °C with a daily 16 h photoperiod for growing seedlings for 10-12 days. At the two-leaf stage, seedlings were uniformly inoculated with a suspension of fresh urediniospores (spores of just harvested or dried spores stored at 4 °C less than two months) in Novec 7100 Engineered Fluid (3M, St. Paul, MN, USA) at the rate of 5 mg/mL using a spray inoculator. Inoculated plants were kept in a dew chamber at 10 °C for 24 h in dark and then placed in a growth chamber set at a diurnal temperature cycle 4–20 °C and a corresponding 8/16 h dark/light cycle.

**Table 1.** *Puccinia striiformis* f. sp. *tritici* races used in seedling tests, including isolates and virulence formulae.

| Race | Isolate | Virulence formula* |
| --- | --- | --- |
| PSTv-1 | 19WA-32 | 1 |
| PSTv-4 | 20WA-19-1 | 1, 6, 9, 17, 27, SP, 76 |
| PSTv-11 | 09-134 | 1, 6, 7, 8, 9, 17, 27, 43, 44, Exp2, 76 |
| PSTv-14 | 17-326 | 1, 6, 7, 8, 9, 17, 27, 43, 44, 85, Exp2, 76 |
| PSTv-17 | 21KS-2-1 | 1, 6, 7, 8, 9, 17, 27, 43, 44, SP, Exp2, 76 |
| PSTv-18 | 11-281 | none |
| PSTv-20 | 93TX-210 | 17 |
| PSTv-21 | 90AR-1 | 6, 17 |
| PSTv-27 | 93-27-Yr10 | 10, 17, 24, 32, 85 |
| PSTv-32 | 21K-41-Yr9 | 6, 7, 8, 9, 44, 85, Exp2 |
| PSTv-37 | 21WA-112 | 6, 7, 8, 9, 17, 27, 43, 44, 85, Exp2 |
| PSTv-44 | 21WA-145-Yr1 | 1, 7, 8, 9, 27, 44, Exp2, 76 |
| PSTv-46 | 21WA-141 | 1, 6, 9, 17, 27, 76 |
| PSTv-48 | 21WA-70-Yr1 | 1, 6, 9, 76 |
| PSTv-52 | 22LA-5 | 6, 7, 8, 9, 17, 27, 43, 44, Exp2 |
| PSTv-64 | 06-19NB | 6, 9 |
| PSTv-119 | 20WA-106 | 6, 7, 8, 17, 27, 43, 44, Exp2 |
| PSTv-120 | 20WA-189-Yr1 | 1, 6, 9, 27, SP, 76 |
| PSTv-220 | 20CA-29 | 1, 6, 7, 8, 9, 17, 27, 43, 44, SP, 85, Exp2, 76 |
| PSTv-326 | 20WA-172-2 | 1, 6, 7, 8, 9, 10, 17, 24, 27, 32, 43, 44, 85, Exp2 |

\*Infection types (0–6 = avirulent, 7–9 = virulent); virulence formula lists *Yr* genes to which the *Pst* race is virulent based on their scores on the 18 genes (*Yr1*, *Yr5*, *Yr6*, *Yr7*, *Yr8*, *Yr9*, *Yr10*, *Yr15*, *Yr17*, *Yr24*, *Yr27*, *Yr32*, *Yr43*, *Yr44*, *YrSP*, *Yr85* (= *YrTr1*), *YrExp2*, and *Yr76* (= *YrTye*) in the *Yr* single-gene differentials (Wan and Chen 2014; Wang et al. 2022).

Stripe rust responses were assessed using the 0–9 infection type (IT) scale described specifically for stripe rust by Line and Qayoum (1992) (Line and Qayoum 1992). IT scores of 0–3 indicated resistant reactions, typically showing no visible symptom or necrosis and chlorosis with no to little sporulation. Scores of 4–6 were classified as intermediate, involving moderate sporulation on chlorotic or necrotic tissues, while scores of 7–9 reflected susceptibility, characterized by heavy sporulation with or without visible necrosis or chlorosis.

### Adult plant phenotyping for stripe rust response

Adult-plant phenotyping for responses to *Pst* was conducted under either natural infection or artificial inoculation conditions during the 2022 and 2023 growing seasons. The spring wheat panel was planted at three locations in Washington State—Pullman, Mount Vernon, and Central Ferry—in 2022 and 2023. The Pullman and Central Ferry sites are in eastern Washington about 50 kilometers apart, and Mount Vernon is in western Washington and about 500 kilometers from Pullman and Central Ferry. The Pullman and Mount Vernon fields are rainfed, and the Central Ferry field is under limited irrigation. The eastern and western Washington sites have different wheat and stripe rust seasons and different *Pst* race compositions due to different environments.

In each year, the spring wheat panel was planted together with other spring wheat nurseries in April at the three locations. The 377 wheat accessions were planted in 50-cm rows with 20 cm between rows with 5-gram seeds in a single row for each line. The susceptible check AvS was planted every 20 rows and surrounding the entire spring wheat experimental field to create uniform stripe rust pressure. The fields were maintained with standard management practices for each region. In both 2022 and 2023, the spring wheat field at Pullman was indirectly inoculated as the adjacent winter wheat field was inoculated in late April with *Pst* urediniospores suspended in in Novec 7100 as described above. The spores used for field inoculation were collected from the same location in 2021 and 2022, respectively, and stored in liquid nitrogen. Stripe rust response was visually scored as IT 0 to 9 as described above, and severity (Sev) was scored using a modified Cobb scale to estimate the percentage of infected leaf area (Peterson et al. 1948). The Mount Vernon field was rated twice at the early jointing stage and the flowering stage, while the Pullman and Central Ferry fields were rated at the flowering to milk stages. Only the stripe rust data of the flowering to milk stage were used in this study to represent the adult-plant response for comparison with the seedling response from the greenhouse tests. Because the stripe rust pressure at Central Ferry was too low to allow reliable data collection in 2023, this location-year environment was discarded. In total, we obtained stripe rust data from five location-year environments that were used for this study.

To obtain a composite measure of resistance, the coefficient of infection (CI) was calculated by multiplying disease severity (%) by the IT score and dividing the result by 10, following the previously described method (Saari and Wilcoxson 2003). This scaling normalizes CI values within a 0 to 90 range, enabling more meaningful comparisons across genotypes and environments.

The survey of field race composition conducted in 2022 and 2023 showed that each location contained a complex mixture of *Pst* isolates, with a few dominant races occurring at high frequency. In 2022, PSTv-37 was the most prevalent race in WA, accounting for approximately 41% of all isolates, followed by PSTv-36 (11%), PSTv-18 (8.5%), and PSTv-39 (7.3%). Several additional races such as PSTv-40, PSTv-41, PSTv-52, PSTv-198, and PSTv-201 were each detected at frequencies between 3–5%, while the remaining races occurred at ≤1%. In 2023, PSTv-37 remained dominant in WA, representing about 61% of all isolates, with PSTv-47 and PSTv-41 (each 7.3%) and PSTv-35 (5.2%) also detected at moderate frequency. All other races were found at ≤1%. Across both years, 13 of the 20 races evaluated in seedling tests were also detected in the field (PSTv-4, PSTv-11, PSTv-14, PSTv-18, PSTv-32, PSTv-37, PSTv-46, PSTv-48, PSTv-52, PSTv-64, PSTv-119, PSTv-120, and PSTv-220), indicating substantial race overlap between the greenhouse seedling tests and field tests. Race composition data were obtained from the Annual Stripe Rust Race Data Reports (https://striperust.wsu.edu/races/data/) (WSU 2022).

### PCA, linkage disequilibrium (LD) and kinship analyses

Principal component analysis (PCA) was performed to assess population structure among the 377 spring wheat accessions. The analysis was conducted using filtered and imputed SNP markers with the “prcomp” function in R (version 4.2.1). The genotype matrix was mean-centered and scaled prior to computation. The proportion of variance explained by each principal component (PC) was calculated by dividing the eigenvalues for each PC by the sum of all PC eigenvalues. Pairwise scatterplots using the first three PCs were generated to visualize clustering and potential subpopulation patterns.

LD was calculated as the squared allele frequency correlation (r²) between SNP pairs using PLINK v1.9 (Purcell et al. 2007), following the method described by (Hill and Robertson 1968). LD decay was analyzed separately for the A, B, and D subgenomes, as well as across the whole genome. Mean r² values were computed for increasing physical distance bins, and LD decay curves were fitted using locally weighted scatterplot smoothing (LOESS) (Cleveland 1979) implemented in the “ggplot2” package. The physical distance at which *r*² dropped below 0.2 was used to define the extent of LD decay.

A genomic relationship matrix (GRM) was computed to estimate pairwise kinship coefficients among accessions. The GRM was calculated using the “A.mat” function in the “sommer” package (Covarrubias-Pazaran 2016), following the previously described method (VanRaden 2008).

### GWAS for seedling stripe rust responses

Association analyses were conducted using the multi-locus mixed model (MLMM) framework as implemented in the GAPIT v3.3 package in R (Segura et al. 2012; Wang and Zhang 2021). Each race was treated as a separate phenotype, enabling the detection of both race-specific and potentially broad-spectrum resistance loci, and improving the resolution and specificity of the mapping results. The MLMM approach iteratively fits fixed effects of significant markers while simultaneously modeling background genetic relatedness as a random effect using GRM (VanRaden 2008). Population structure was corrected by incorporating PCs derived from PCA on the genotype matrix. The number of PCs in the model was optimized separately for each race by visually inspecting quantile–quantile (Q–Q) plots. Significance thresholds for marker–trait associations were determined using the false discovery rate (FDR) (Benjamini and Hochberg 1995), with a cut-off of FDR ≤ 0.05. Following model fitting, results were summarized across races to identify both race-specific and overlapping resistance loci. To facilitate the visualization of genome-wide marker–trait associations for each race, Manhattan plots were generated using the “ggplot2” R package (Wickham 2016), highlighting the chromosomal distribution and statistical significance of SNPs across the wheat genome. Peaks corresponding to significant associations were further examined for their proximity to known resistance loci and for potential pleiotropic effects across races.

### GWAS for adult-plant stripe rust responses

GWAS was carried out using phenotypic data from 15 trait–environment combinations. These combinations included IT, Sev, and CI scores collected from field trials conducted in Pullman, Mount Vernon, and Central Ferry, Washington, during the 2022 and 2023 growing seasons. Each unique trait-location-year combination was treated as an individual phenotype, enabling the identification of both environment-specific and stable QTLs associated with APR. As with ASR, the association analysis was conducted using the MLMM model (Segura et al. 2012; Wang and Zhang 2021) following the same procedures of population structure and relatedness correction. To determine the optimal number of PCs, Q–Q plots were evaluated for each trait–environment combination, ensuring that the model was neither too permissive (leading to p-value inflation) nor too conservative (leading to reduced power). This optimization step was crucial given the environmental variability and trait differences across locations and years. Regions with multiple significant SNPs, consistent signals across environments and not overlapping with ASR loci were considered as potential APR loci.

### LD with known *Yr* genes and QTLs

To detect overlap between GWAS-identified loci and previously reported stripe rust resistance regions, a targeted linkage disequilibrium (LD) analysis was conducted using a comprehensive reference set of 1,144 known loci. This set combined previously curated *354* Yr genes and QTLs (Tong et al. 2024), originally mapped to RefSeq v2.1 (Zhu et al. 2021), and 790 additional resistance loci reported by Wu et al. (2025), which were already aligned to the IWGSC RefSeq v1.0 reference genome (The International Wheat Genome Sequencing Consortium (IWGSC) 2018). To ensure coordinate compatibility with our SNP dataset, physical positions from Tong et al. (2024) were converted to RefSeq v1.0 coordinates, while no conversion was required for the loci from Wu et al. (2025).

For each known resistance locus, a ±10 Mb physical window was defined, and all SNPs within these intervals were extracted from the filtered genotype matrix. Pairwise LD (r²) was calculated between significant GWAS SNPs and the SNPs located within these predefined windows using PLINK v1.9 (Purcell et al. 2007). SNP pairs with r² ≥ 0.6 were retained as evidence of potential co-localization. Loci meeting this threshold were classified as previously reported or known, while those showing no evidence of overlap or LD with any reference loci were retained as putatively novel associations.

## Results

### Seedling responses to stripe rust

The 377 spring wheat accessions displayed a wide range of responses to infection with 2 *Pst* races in the seedling stage, with IT scores ranging from 0 (immune) to 9 (fully susceptible) (Fig. 1a, Table S2). Based on a resistance threshold of IT ≤ 6, only two races were avirulent in more than 65% of accessions—PSTv-18 (83.02%) and PSTv-64 (68.97%). For the majority of the races (18 out of 20), over 50% of accessions showed susceptible reactions (IT ≥ 7), indicating the rarity of effective ASR alleles within the panel. The most virulent races were PSTv-37 and PSTv-326, for which only 2.65% and 1.86% of accessions, respectively, exhibited resistance. Overall, the panel exhibited race-specific variation, with substantial differences in resistance frequencies across races.

**Fig. 1.**
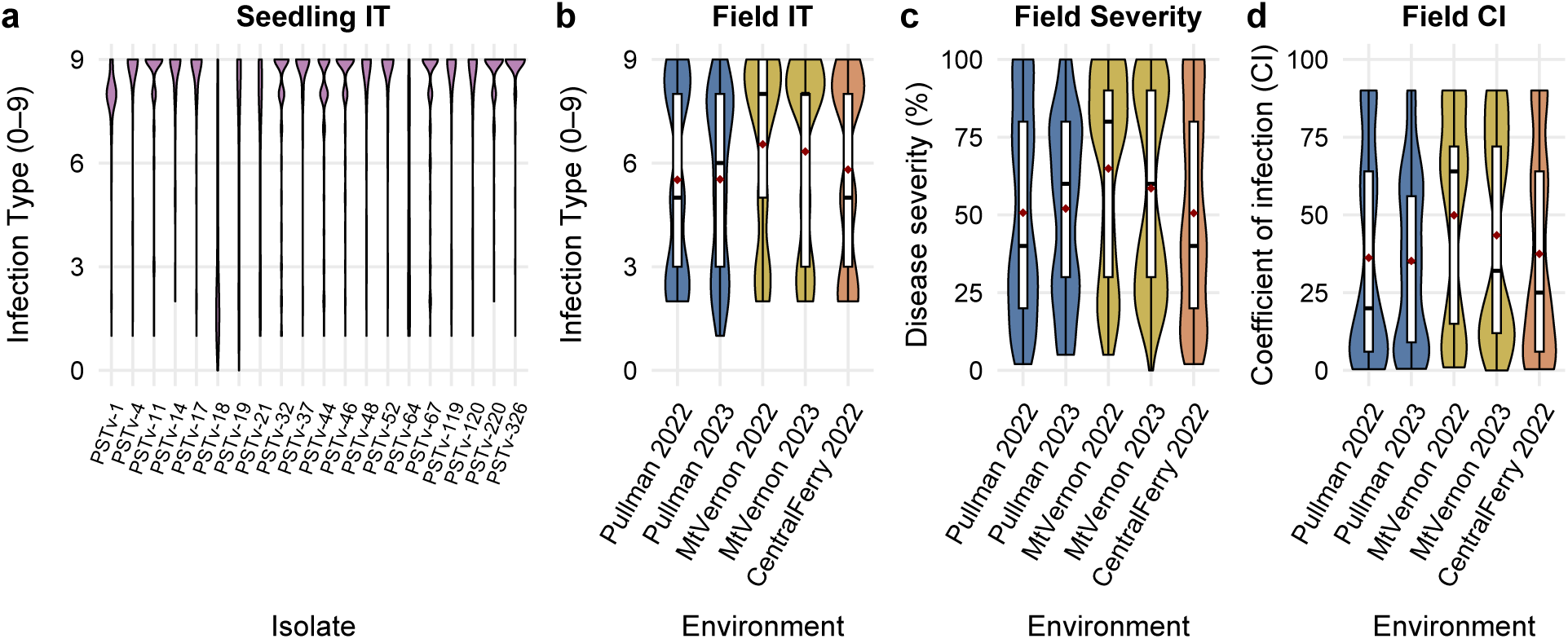
Distribution of stripe rust response scores in the diverse wheat panel. **(a)** Violin density plots show the distribution of seedling infection type (IT) scores collected for 377 wheat lines under controlled greenhouse conditions. **(b - d)** Violin density plots with embedded boxplots show the distribution of IT scores (b), disease severity scores (c) and coefficient of infection (CI) (d) evaluated at the adult-plant stage in field trials across three locations in 2022 (Pullman, Mount Vernon, and Central Ferry) and two locations in 2023 (Pullman and Mount Vernon).

A total of 50 accessions (13.3%) were fully susceptible, exhibiting IT scores ≥ 7 against every race tested (Table S2). The remaining accessions displayed variable responses across isolates, indicative of the presence of race-specific resistance genes with intermediate to major effects on disease progression. Eleven spring wheat lines from the diversity panel exhibited resistance to at least 75% of the 20 *Pst* races evaluated at the seedling stage (Table S2). Notably, lines such as GF104 (CItr 15089), GF161 (PI 189794), GF225 (PI 207115), GF231 (PI 366981), and GF257 (PI 137758) showed resistance to 18 or more of the 20 *Pst* races, including highly virulent pathotypes PSTv-37, PSTv-52, PSTv-220, and PSTv-326. Among them, GF104 and GF225 were resistant to all 20 races and also remained resistant at the adult plant stage. These genotypes represent valuable sources of ASR for breeding programs focused on enhancing durable and broad-spectrum resistance to stripe rust. These findings highlight the diversity of ASR genes within the panel and emphasize the importance of screening diverse germplasm to identify rare accessions with broad-spectrum responses for use in resistance breeding programs.

### Adult-plant responses to stripe rust

The stripe rust pressure from either natural infection (Central Ferry and Mount Vernon) or artificial inoculation (Pullman) was high across all five environments, as shown by the consistent IT 9 and 100% severity of the susceptible check ‘Avocet S’ in the five location-year environments (Fig. 1b., Table S3, Table S4). The uniformity of ‘Avocet S’ responses across the location-year environments demonstrates that the stripe rust phenotype data were suitable for GWAS analysis. Each trial (location-year combination) was analyzed independently for IT, Sev, and CI, providing insights into the genetic basis of and environmental effect on ASR and/or APR observed at the adult-plant stage in the fields.

In 2022, mean IT values ranged from 5.52 ± 2.71 in Pullman to 6.54 ± 2.55 in Mount Vernon, while CI ranged from 36.23 ± 32.36 to 49.88 ± 32.33, respectively (Table S5). Central Ferry, evaluated only in 2022, had an intermediate IT profile (IT = 5.81 ± 2.69; CI = 37.47 ± 31.97), with fewer highly susceptible lines. In 2023, stripe rust pressure remained stable, with Pullman and Mount Vernon showing mean ITs of 5.53 ± 2.51 and 6.33 ± 2.66, respectively. IT score distributions (Fig. 1b) further illustrated these differences: Pullman displayed broader phenotypic variability, indicating greater genotypic diversity in resistance levels, while Mount Vernon exhibited more compressed distributions, with a larger proportion of susceptible responses. The percentage of accessions classified as resistant (IT ≤ 3) ranged from 22.7% (Mount Vernon 2022) to 29.6% (Central Ferry 2022) and increased slightly in 2023 to 26.8% (Mount Vernon) and 29.3% (Pullman), highlighting both spatial and temporal variation in adult-plant resistance expression.

Severity and CI distributions (Fig. 1c, d) followed a similar pattern: Pullman showed a wider range of values and a larger proportion of moderately resistant lines, while Mount Vernon exhibited higher mean severity and CI scores with more uniform, susceptible responses. The higher mean severity and CI values observed at Mount Vernon likely reflect the local race composition, which was dominated by highly virulent races such as PSTv-37, along with PSTv-36, PSTv-18, and PSTv-39 in 2022. Similarly, in 2023, PSTv-37 remained predominant, accompanied by PSTv-47, PSTv-41, and PSTv-35. The prevalence of these aggressive races likely contributed to the higher mean IT, Sev, and CI values and the reduced variability observed at this location.

Among the tested lines, 84 were resistant (IT ≤ 3) or intermediately resistant (IT ≤ 6) in at least one field environment. Of these, 40 accessions were classified as highly resistant in at least one trial. Notably, GF75 (PI 351907) and GF358 (PI 272377), which displayed complete susceptibility at the seedling stage (IT ≥ 7 to all 20 *Pst* isolates), exhibited consistent adult-plant resistance across both years, with field IT scores reduced to 0–3 and lower disease severities, suggesting the presence of non-race specific APR. Conversely, GF104 (CItr 15089) and GF225 (PI 207115) showed resistance to all races in the seedling tests and resistant / intermediate resistant responses across most field environments, indicative of broad-spectrum ASR.

Correlation analysis among environments revealed moderate to high consistency in disease response traits, with the strong correlations (~0.7) observed between Pullman 2022 and Pullman 2023 as well as between Mount Vernon 2022 and Mount Vernon 2023 across all three metrics (IT, Sev, and CI), suggesting stable field resistance at these two sites. These findings support the importance of multi-environment and multi-year evaluations for identifying durable APR and effective ASR sources suitable for incorporation into resistance breeding programs.

### GWAS for seedling stripe rust responses

For GWAS, 625,579 markers were assigned to the A genome, 773,265 to the B genome, and 123,014 to the D genome. Principal component analysis showed that PC1, PC2 and PC3 explained 9.06%, 6.24%, and 3.47% of the total genetic variance, respectively, with the first ten components cumulatively accounting for 30.98% of the variation. This moderate level of population structure suggests the lack of strong underlying genetic stratification within the panel. Genome-wide LD analysis revealed variation in LD decay rates across subgenomes, with r² values dropping below 0.2 around 950–1,000 kb in the A and B genomes, while the D genome exhibited extended LD, maintaining r² ≥ 0.2 up to approximately 1,300 kb. These levels of LD suggest that the diversity panel should provide sufficiently high levels of resolution for mapping genes to small genomic intervals.

Genome-wide association mapping ASR response to 20 distinct *Pst* isolates revealed 68 non-redundant lead SNPs identified by grouping all significant marker-trait associations based on LD threshold of *r*² ≥ 0.6 in 10 Mb window (Table S6, Fig. 2a, c). These ASR loci were unevenly distributed across the three subgenomes, with the A, B and D genomes harboring 22, 41, and 5 SNP-trait associations, respectively. Notably, chromosome 3B emerged as a resistance hotspot, harboring 11 SNPs in total, followed by chromosome 1A, which harbored seven significant SNPs. Across the entire GWAS dataset, seven SNPs were significantly associated with IT scores for up to four distinct *Pst* isolates (Table 2), suggesting that certain genomic regions may provide broader-spectrum ASR across the wheat panel. While most SNPs were race-specific and rare, this subset of multi-isolate SNPs may represent promising targets for functional validation and introgression into breeding lines. The number of significant SNPs per race varied widely (Fig. 2b), with PSTv-36 and PSTv-1 contributing the most associations (9 and 8 SNPs, respectively), while several races showed only one or two associations.

**Fig. 2.**
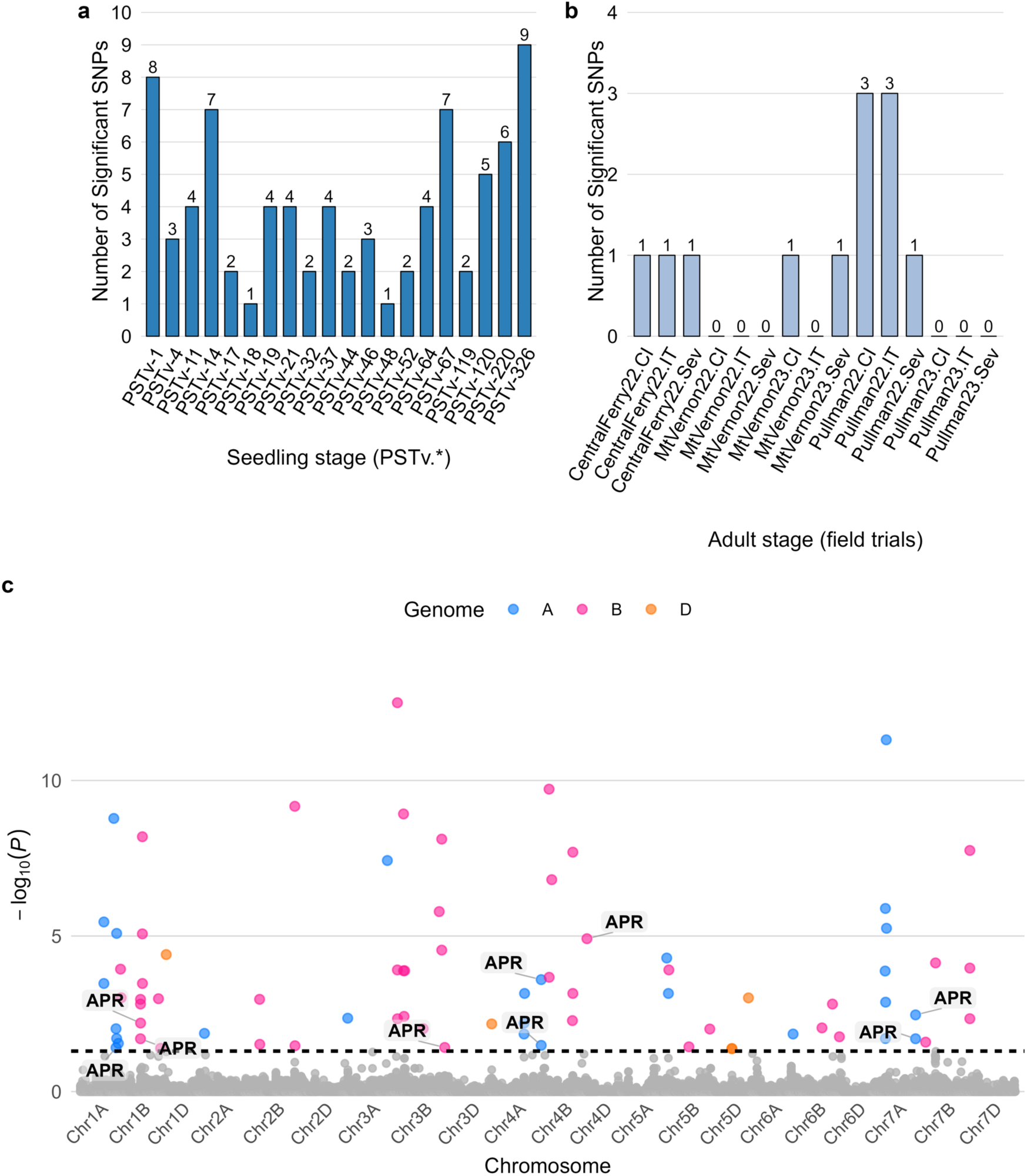
Summary of GWAS results for stripe rust resistance in the greenhouse seedling screens and field trials of adult stage plants. **(a)** Number of significant SNPs detected in the seedling tests with 20 *Puccinia striiformis* f. sp. *tritici* races. **(b)** Number of significant SNPs identified in filed trials conducted at different locations and scored for coefficient of infection (CI), disease severity (Sev), or infection type (IT). **(c)** Manhattan plot showing the genomic positions and FDR-corrected significance levels (−log₁₀P) for SNPs associated with resistance across seedling and adult stage trials. SNPs are color-coded by genome (A, B, or D). The dashed line represents the significance threshold (α = 0.05). Significant SNPs identified in the field trials at the adult plant stage are labelled “APR”.

**Table 2.**
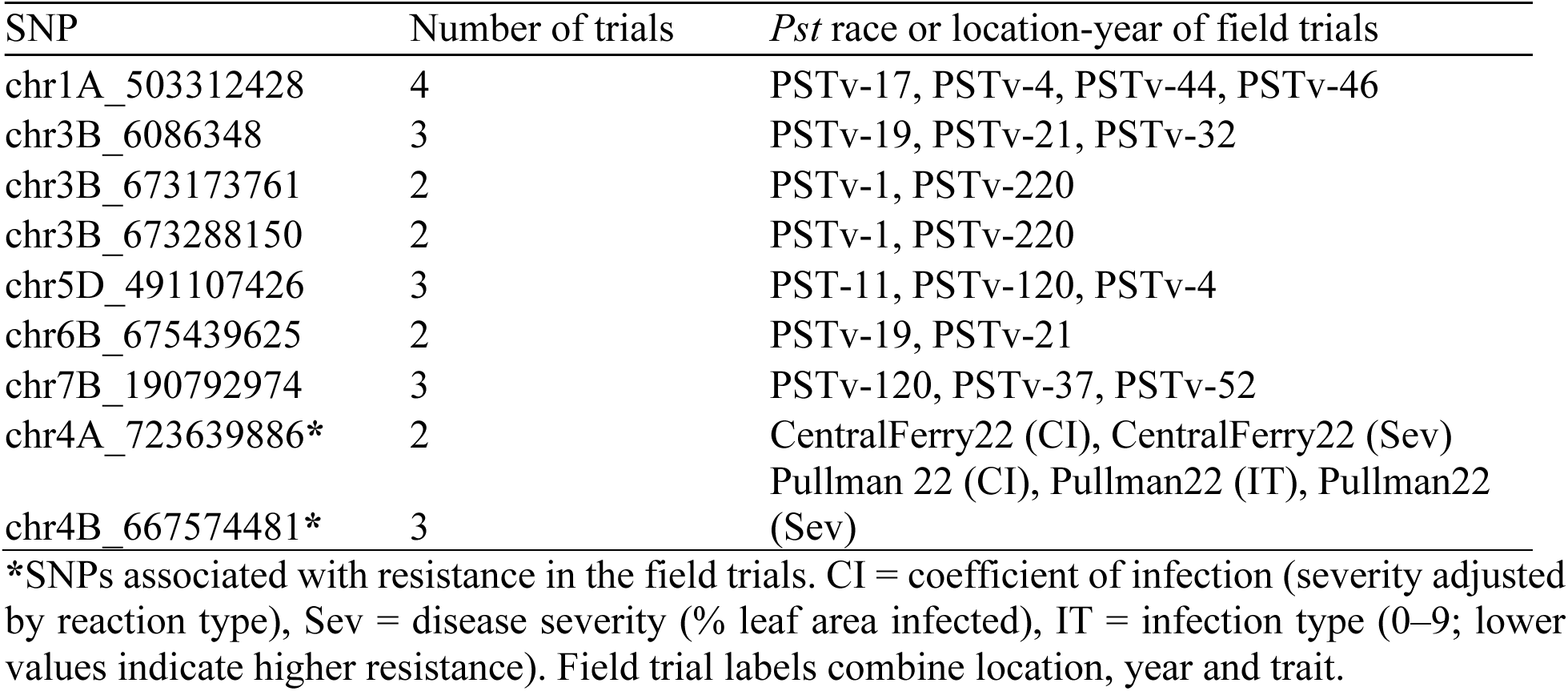
Significant SNPs detected in more than one trial for seedling and adult stage resistance to *Puccinia striiformis* f. sp. *tritici*.

### GWAS for adult-plant stripe rust responses

The GWAS for stripe rust resistance in the field trials was based on 15 distinct trait–year-environment combinations, including three locations (Pullman, Central Ferry and Mt. Vernon) and two growing seasons (2022 and 2023). All these locations showed extensive variation in the composition of field *Pst* populations in WA in 2022 and 2023, with each locations including multiple races (https://striperust.wsu.edu/races/data/). The Central Ferry location data from the 2023 field trial was excluded due to low stripe rust pressure (Table S6).

In total, nine non-redundant SNPs were significantly associated with IT, Sev, and CI (Fig. 2b, c). These SNPs were distributed among six chromosomes, including 1A, 1B, 3B, 4A, 4B, and 7A (Fig. 2c). The significant SNP detected for IT, CI, and Sev traits on chromosome 4B (chr4B_667574481) was detected only during the Pullman 2022 trial. Likewise, two significant SNPs, one associated with CI and Sev (chr4A_723639886) and another with IT (chr4A_723810464) were detected only during the Central Ferry 2022 trial. These results suggest the presence of environment-specific SNPs likely associated with variation in the race composition of local Pst populations or with other genotype-by-environment interactions that influence disease development. No overlap was detected between these nine SNPs and the ARS loci mapped in greenhouse seedling screens. The association of these SNPs with non-race-specific resistance responses to complex field mixtures of Pst races suggests that they likely represent APR loci.

### Frequency of known *Yr* resistance genes

To assess the presence of previously characterized *Yr* resistance loci in our diversity panel, a set of five major resistance genes, including *Yr5*, *Yr15*, *Yr36*, *Lr34*/*Yr18*, and *Lr46*/*Yr29*, was evaluated using functional KASP markers. The results revealed a markedly low occurrence of these major resistant alleles in the panel. Specifically, the resistance allele for *Yr5* was detected in only 12 accessions (3.2%), while *Yr15* was completely absent in the panel. These two genes confer broad-spectrum ASR and remain effective against all *Pst* races used in this study. Their absence from the panel likely reflects their relatively recent discovery and targeted introgression into modern elite cultivars, particularly in structured breeding programs, rather than widespread distribution in traditional or landrace germplasm (Table S8).

Among the APR genes, only *Lr46*/*Yr29* was present at a moderate frequency, identified in 147 accessions (39%), while *Yr36* was found in just 10 accessions (2.7%). The widely studied slow-rusting gene *Lr34*/*Yr18,* known for conferring durable resistance in many modern cultivars, was completely absent in the panel. The near-complete lack of *Yr36* and *Lr34*/*Yr18* suggests that these APR sources have not been extensively deployed in the geographic or temporal scope represented by the selected lines (Table S8).

Taken together, the absence or low frequency of these key *Yr* resistance alleles suggests that our diversity panel captures stripe rust resistance alleles that have not been the target of strong selection. This underscores both a gap in the historical deployment of these loci and a potential opportunity for breeders to exploit this diversity for novel or complementary resistance sources that may function independently of the well-characterized genes tested in this study.

### Overlap of mapped loci with the previously reported *Yr* loci

To investigate whether the SNPs identified through GWAS correspond to previously reported *Yr* loci (Tong et al. 2024; Wu et al. 2025), a targeted LD-based comparison was performed. Of the 77 significant SNPs identified in our study, 33 (36%) showed strong LD (*r*² ≥ 0.6) with 43 known *Yr* genes or QTLs (Fig. 3). Among these, nine SNPs were in perfect LD (*r*² = 1) with at least one known *Yr* locus. This considerable overlap underscores the robustness of the GWAS findings and suggests that approximately 36% of detected associations are located within genomic regions harboring previously reported *Yr* loci (Fig. 3; Table S9).

**Fig. 3.**
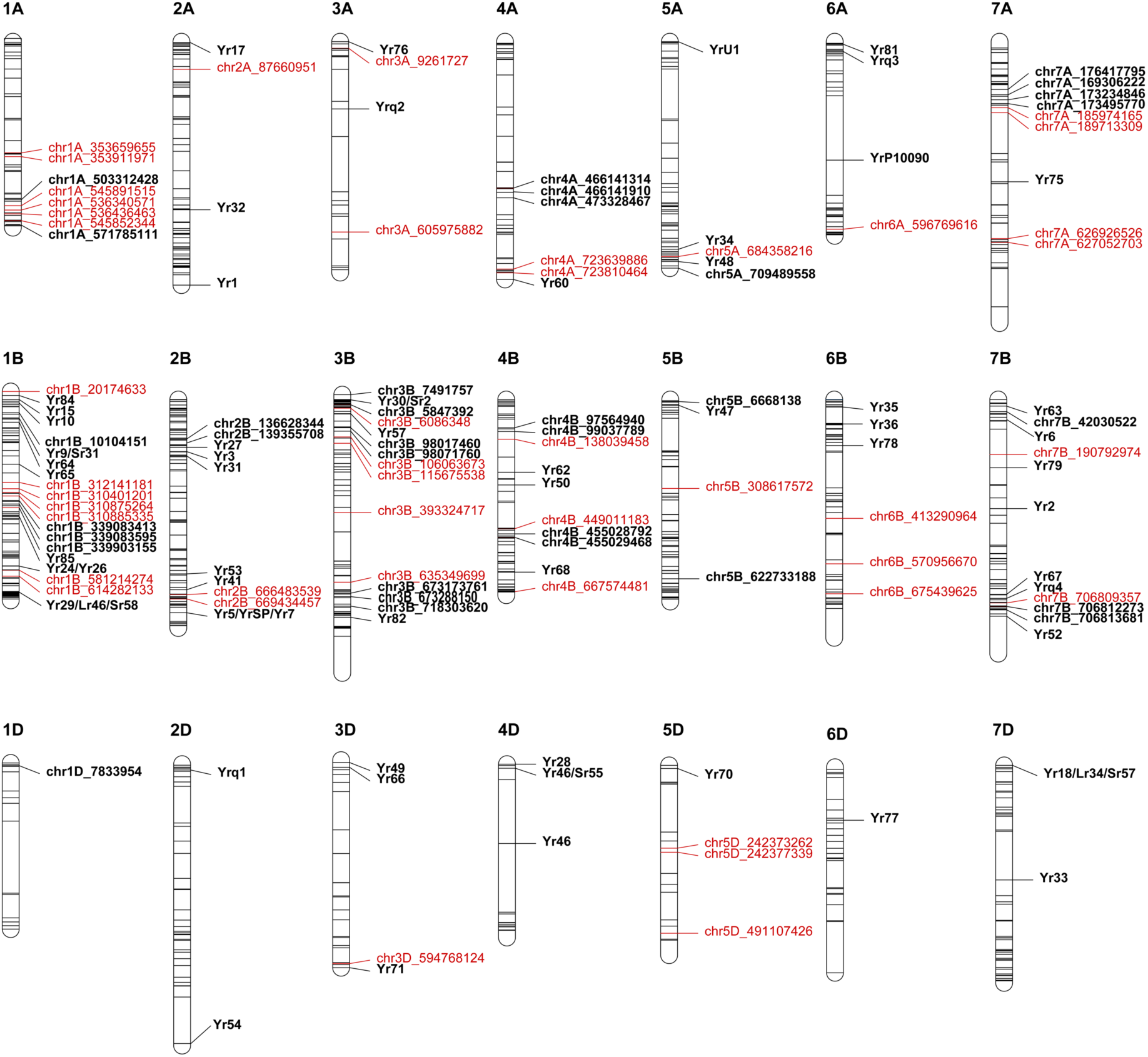
Physical map of wheat *Yr* loci across wheat chromosomes. Black markers show locations of the previously reported *Yr* genes and QTLs, whereas red markers show significant SNPs identified in our study. Black markers with locus names correspond to the known *Yr* resistance loci that overlap with loci detected in our study. Red markers with locus names denote novel loci that do not coincide with the known *Yr* loci.

The chromosomal distribution of overlapping loci shows that chromosome 3B harbored the highest number of known resistance loci (n = 13), followed by chromosomes 2B (n = 5) and 5B (n = 4). Of particular interest, a single region on chromosome 3B displayed a high density of overlapping loci, suggesting the existence of a haplotype block enriched for stripe rust resistance. For example, chr3B_5847392 was found to be in strong LD (r² = ~0.95) with 13 known resistance loci, including the multi-pathogen resistance gene complex *Sr2*/*Lr27*/*Yr30*, highlighting this locus as a high-value genomic region with potential broad-spectrum effectiveness.

We detected one APR locus located on chromosome 3B (chr3B_718303620) that overlapped with the previously mapped stripe rust resistance locus *QYr.agis104* (Cheng et al. 2024). There was also one SNP, chr1B_614282133, that mapped 47 Mb away from *Yr29*/*Lr46* (William et al. 2003) located at 661 Mb. Combined with the results of KASP genotyping based for known APR genes, these results suggest that the majority of APR loci detected in our study are novel.

Among the overlapping loci, two well-characterized *Yr* genes, *Yr30* and *Yr47*, were identified within genomic regions exhibiting strong LD with GWAS SNPs. *Yr30*, a component of the *Sr2*/*Lr27*/*Yr30* multi-pathogen resistance locus, was co-localized with chr3B_5847392. Likewise, *Yr47* was detected in perfect LD (r² = 1) with chr5B_6668138, a GWAS-associated SNP also co-localized with three major QTLs: *QYr.eu30*, *QYr.inra-5BL.2*, and *QYr.uga-5B*. Together, the identification of *Yr30* and *Yr47* in regions of strong LD with GWAS-significant markers not only reinforces the biological significance of these loci but also underscores their utility as strategic targets in breeding programs focused on developing cultivars with enhanced and sustained resistance.

### Wheat improvement had distinct impact on ASR and APR alleles

Pyramiding ASR and APR genes increases resistance to wheat rusts (Zhang et al. 2019; Wang et al. 2023; Wu et al. 2025). While our results, in general, are consistent with these observations for *Yr* loci detected in both seedling and adult stage screens (Fig. 4a), we also found that the phenotypic effects of APR and ASR accumulation have distinct dynamic. Increase in the number of ASR alleles had little effect on mean IT of the majority of wheat lines in the panel. This is likely associated with the fact that the majority of *Yr* genes mapped in the seedling screens are race-specific and have limited contribution to mean IT calculated from data obtained using 20 *Pst* races. Only limited fraction of wheat lines showed mean IT ≤ 6 (Fig. 4a), including GF225 and GF104 lines, which were resistant against all 20 tested *Pst* isolates. Contrary to ASR loci, combinations of 4-5 APR alleles in many wheat lines was sufficient to provide good levels of resistance (IT ≤ 6) in the field trials. The three-fold higher frequency of resistant alleles at APR loci compared to those at ASR loci (Mann–Whitney U-test p-value = 2.6 × 10^−110^) could have also facilitated accumulation of multiple APR alleles in some wheat lines (Fig. 4b).

**Fig. 4.**
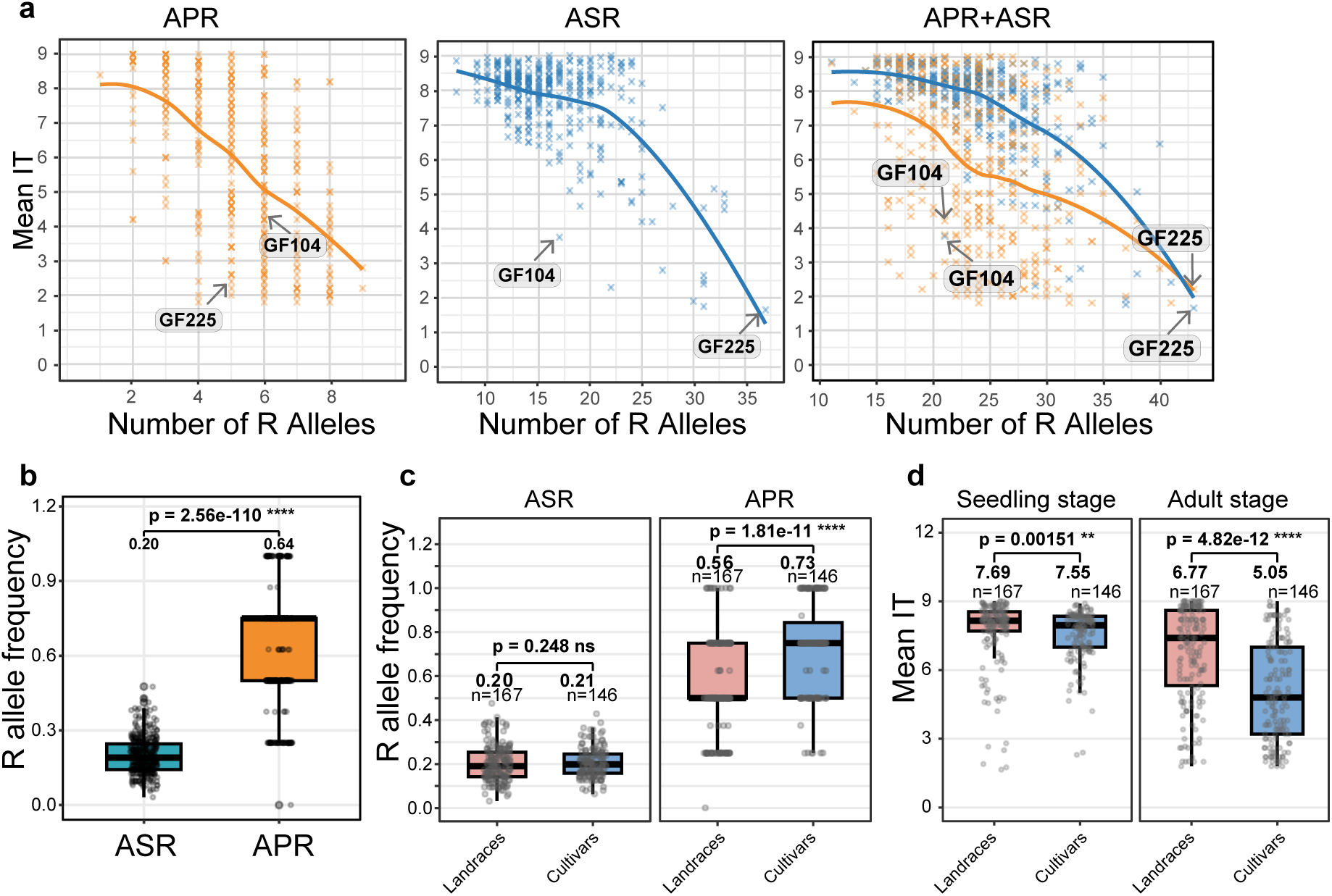
Distribution of resistance (R) gene loci across populations and among wheat lines. (a) The X-axis shows the number of R alleles per line. Y-axis shows the mean infection type (IT) scores based on data from 20 *Pst* races for ASR and field infection trials for APR. Both adult plant and seedling stage screening data were combined for assessing the impact of total number of APR and ASR R alleles on IT. The data collected at the adult and seedling stages is shown as orange and blue crosses, respectively. (b) Comparison of R allele frequencies between ASR and APR loci. A nonparametric Mann–Whitney U-test was applied to compare allele frequencies. (c) Comparison of ASR and APR allele frequencies between landraces and cultivars. Within each growth stage, group differences were tested using a one-sided Mann–Whitney U-test. (d) Comparison of IT collected at the seedling and adult plant stages between landraces and cultivars. Data were grouped into Landraces and Cultivars (cultivars and breeding selections), excluding genetic resources. Mean IT values and sample sizes (n) are indicated above each box, and horizontal brackets denote p-values from one-sided Mann–Whitney U-test. Significance levels: **** p < 0.0001; *** p < 0.001; ** p < 0.01; * p < 0.05; ns = not significant.

The differences in the accumulation of resistant alleles between APR and ASR loci could be associated with differences in the historic selection that acted on these loci. Selection for ASR, which are mostly represented by race-specific *Yr* genes, varies across time and space and largely driven by ‘boom and bust cycles’ of wheat-*Pst* interaction. The *Yr* loci detected in field trials under natural infection conditions, which include mixes of *Pst* isolates varying from year-to-year in composition, likely include few all-stage race-specific resistance genes and mostly APR loci that have more quantitative nonrace-specific effects. If this assumption is correct, we may expect that selection associated with wheat improvement preferentially targeted APR alleles rather than ASR alleles. To test this hypothesis, we compared the changes in resistant allele frequency for APR and ASR loci between landraces and more modern improved wheat cultivars, including breeding material and cultivars. Consistent with our assumption, we observed no changes in resistant allele frequency for AST between landraces and cultivars (one-sided Mann–Whitney U-test, p-value = 0.25), whereas resistant alleles at APR loci showed 30.3% increase in cultivars compared to landraces (one-sided Mann–Whitney U-test, p-value = 1.8 × 10^−11^) (Fig. 4c). The changes in IT mirrored the results of resistant allele frequency comparison; transition from landraces to cultivars was associated with only 1.8% reduction in IT for seedling stage resistance (one-sided Mann–Whitney U-test, p-value = 1.5 × 10^−3^), whereas IT at adult stage reduced by 25.5% (one-sided Mann–Whitney U-test, p-value = 4.8 × 10^−12^).

## Discussion

The continued evolution of *Pst* has rendered many resistance genes deployed in wheat cultivars ineffective, increasing the urgency to identify novel and durable sources of resistance (Wan and Chen 2014; Schwessinger 2016; Shahinnia et al. 2022; Riella et al. 2024; Davis et al. 2025). This study used a globally representative panel of wheat lines, including landraces, historical cultivars, and breeding lines, to evaluate resistance to stripe rust. The panel spans more than a century of wheat development and captures geographic and genetic diversity from key centers of domestication and adaptation (He et al. 2022). This region-specific selection led to accumulation of a broad spectrum of resistance alleles, including those that are underrepresented in modern breeding programs (Adhikari et al. 2012; Zegeye et al. 2014). Using a diverse set of 20 *Pst* isolates we discovered a wide range of infection types, revealing considerable variation in resistance responses and the potential to uncover both established and novel resistance loci.

ASR genes effective against multiple stripe rust isolates, especially *Yr5* and *Yr15*, are broadly used in developing resistant wheat varieties (Yaniv et al. 2015; Klymiuk et al. 2018; Marchal et al. 2018) and represent valuable targets for gene discovery efforts. Five lines from our diversity panel (GF104, GF161, GF225, GF23, and GF257) showed resistance to 18 or more of the 20 *Pst* races, including the highly virulent races PSTv-37, PSTv-52, PSTv-220, and PSTv-326. Among these lines, GF104 (CItr 15089) and GF225 (PI 207115) were resistant to all 20 races, with both lines being resistant at the adult plant stage too. Diagnostic marker analysis revealed that none of the lines in our diversity panel carried well-characterized ASR genes *Yr5* (Marchal et al. 2018) and *Yr15* (Klymiuk et al. 2018), indicating that other, potentially novel or under-characterized ASR loci may contribute to the observed resistance in GF104 and GF225. In light of recent reports that resistance conferred by Yr5 and Yr15 has been broken (Tekin et al. 2021; Davis et al. 2025), identifying wheat lines that remain resistant to a broad range of Pst races provides potentially novel sources of ASR genes for breeding durable stripe rust resistance.

Among the APR genes screened using KASP markers in our panel, three well-characterized loci, *Yr29*/*Lr46*, *Yr18*/*Lr34*, and *Yr36* (William et al. 2003; Lagudah et al. 2009; Fu et al. 2009) were specifically targeted due to their roles in providing durable, non-race-specific resistance to stripe rust. These genes function through partial resistance mechanisms and are often more effective when combined with other *Yr* genes, showing additive effects in gene pyramiding strategies (Singh et al. 2016; Wang et al. 2023). They are also associated with pleiotropic resistance to multiple pathogens, including leaf, stem and stripe rust (Krattinger et al. 2009; Lagudah 2011). In our panel, only *Yr29*/*Lr46* was present at high frequency (39%), while *Yr36* was present at low frequency (2.7%) and *Yr18*/*Lr34* was not detected. The lack of overlap between the *Yr* loci detected in our study and these major APR genes indicate that variation in resistance at the adult plant stage is largely driven by alternative, potentially novel sources of APR in the panel. For example, two lines in the panel, GF75 and GF358, were susceptible to all 20 *Pst* races at the seedling stage but consistently exhibited resistance or intermediate resistance across all adult plant field trials. GF75 tested positive for *Yr29*/*Lr46*, while GF358 lacked all three APR markers screened in this study (*Yr18*/*Lr34*, *Yr29*/*Lr46*, and *Yr36*). These examples confirm that lines in our diversity panel harbor the APR sources that have yet to be fully characterized.

In spite of extensive prior mapping efforts focused on discovery of the *Yr* resistance genes in diverse germplasm (Tong et al. 2024; Wu et al. 2025), almost 60% of the *Yr* loci mapped in our study showed no linkage with the previously mapped genes. Some previous GWAS studies, particularly those utilizing narrower genetic panels or fewer pathogen races, reported higher co-localization rates with known resistance loci (Maccaferri et al. 2015; Shahinnia et al. 2022). This relatively lower match rate may reflect the broader genetic diversity and greater environmental and pathogen variation encompassed in our panels of wheat lines and *Pst* isolates, leading to the discovery of novel and previously uncharacterized associations.

One of the interesting aspects of our data are the significant differences in population R allele frequency observed for *Yr* genes detected in the seedling and adult-plant screens. These differences are likely associated with the differences in the expression of disease resistance traits controlled by the mapped *Yr* genes. The ASR *Yr* genes are usually associated with race-specific strong dominant resistance expressed at all stages of development and could be easily selected during breeding process (Chen 2013; Wang et al. 2023). The strong phenotypic effects expressed by this class of resistance genes also apply strong selection pressure on pathogen populations leading to the origin of new races that overcome resistance (Brown 2015; Riella et al. 2024). The shifts in the composition of pathogen populations refocus local breeding efforts on other strong effect resistance genes that confer resistance against newly emerged pathogen races. As a result, most ASR genes never reach high frequency in diverse wheat panels, which is consistent with our data. On the contrary, many APR *Yr* genes express moderate to weak race non-specific resistance and likely impose weaker selection pressure on pathogen populations, preventing quick evolution of new virulent races. At the times when major resistance genes became non-effective due to evolution of virulent races, it is likely that selection would favor genotypes that carry the race non-specific genes with weaker effects. In addition, these loci potentially enhance resistance when combined with ASR genes (Zhang et al. 2019; Wang et al. 2023). Both factors can contribute to consistent long-term selection pressure on quantitative disease resistance loci and drive their frequency to higher levels in more modern germplasm, as was observed in our diversity panel.

In conclusion, this study demonstrates the value of continued screening of diverse global germplasm using diverse collections of pathogen races for discovering novel sources of resistance for breeding. Variation in the representation of ASR and APR loci in various diversity panels combined with extensive variation at avirulence loci in the populations of pathogens (Olivera et al. 2019; Guo et al. 2022; Riella et al. 2024) offer unique opportunities for uncovering the genetic makeup of interaction between wheat and its pathogens and identifying novel loci to protect wheat production from disease outbreaks. Using a panel composed of landraces, historical cultivars, and breeding lines, we identified 77 *Yr* loci conferring ASR or APR. A total of 44 *Yr* loci were potentially novel sources of resistance that warrant further investigation. Importantly, we identify two accessions, GF104 (CItr 15089) and GF225 (PI 207115), that exhibited resistance to all 20 *Pst* races yet lacked diagnostic markers for the broadly effective ASR genes *Yr5* and *Yr15*, suggesting that their resistance is conferred by previously uncharacterized ASR alleles. Overall, these findings not only confirm the value of known *Yr* loci but also expand the catalog of candidate loci available for marker-assisted selection. Together, our results provide actionable targets for diversifying the genetic base of resistance and advancing the development of wheat cultivars with broad-spectrum, durable resistance against evolving *Pst* populations.

## Statements and Declarations

### Data Availability

Phenotyping data collected in this study is provided as supplementary information. The germplasm is available through the USDA-ARS Germplasm Resources Information Network (GRIN).

## Acknowledgements

We would like to thank Kent Evans for helping plant the wheat panel in fields. We would like to thank Kent Evans for helping plant the wheat panel in fields. The mention of trade names or commercial products in this publication is solely to provide specific information and does not imply recommendation or endorsement by the United States Department of Agriculture. The USDA is an equal opportunity provider and employer.

## Authors contribution

BS – GWAS analyses, genotyping and preparation of manuscript draft; MW – phenotyping for stripe rust resistance; JDZ – GWAS analyses; XX – genotyping known stripe rusts resistance loci; GB - genotyping known stripe rusts resistance loci; AA – sequence-based genotyping of a diversity panel; XC - phenotyping for stripe rust resistance; ED – funding acquisition, conceived project, data analysis, manuscript writing. All authors reviewed the manuscript.

